# Maximizing the fidelity of a photovoltaic subretinal prosthesis for human patients

**DOI:** 10.1101/2025.03.31.646451

**Authors:** Nathan Jensen, Keith Ly, Anna Kochnev Goldstein, Quentin Devaud, Daniel Palanker

## Abstract

**Objective:** PRIMA subretinal implants provide prosthetic vision to patients blinded by age-related macular degeneration, with acuity closely matching the sampling limit of the pixel pitch: a single 100µm pixel per line of a letter corresponds to 20/420 acuity. Decreasing the pixel size in the same flat geometry is difficult due to the constrained electric field, especially considering a 40µm thick debris layer separating the implant from the target neurons. Here we optimize the electrode design to help overcome such limitations.

**Approach:** An end-to-end modeling pipeline combines the retinal photovoltaic implant simulator (RPSim) based on the Xyce circuit simulator with an interface to COMSOL Multiphysics for electric field modelling. It was used to generate and characterize implants in an open-loop sampling-based optimization. Implant performance was evaluated with respect to voltage drop across bipolar cells (representing the stimulation strength), pattern contrast, and neural selectivity.

**Main Results:** The highest selectivity in stimulation of bipolar cells was achieved with arrays having active electrodes on pillars and return electrodes connected in a mesh surrounding the photovoltaic pixels in the array. Such a design, even with pixels down to 20µm, provides stimulation strength exceeding, and contrast similar to that of flat 100µm PRIMA pixels.

**Significance:** Using a novel 3-D electrode design, the pitch of the photovoltaic array can be decreased to 20µm, while providing performance that exceeds the flat 100µm PRIMA pixels. In humans, 20µm resolution on the retina corresponds to a visual acuity of 20/80 – a five times improvement compared to the current clinical device.

## I. Introduction

The retinal prothesis system PRIMA provides central vision in patients impaired by atrophic age-related macular degeneration [1]. It includes a wireless photovoltaic subretinal implant and a pair of augmented reality (AR) glasses equipped with a video camera. Images captured by the camera are processed and projected onto the implant using pulsed light. The implant itself is a 2×2mm multielectrode array (MEA) with hexagonal photovoltaic pixels (Fig. 1(a)) that convert this light into electric current to stimulate the nearby retinal neurons in the inner nuclear layer (INL), primarily the bipolar cells (BCs). To avoid stimulation of the healthy peripheral retina, the glasses use an invisible near-infrared (NIR, 880 nm) wavelength.

**Fig. 1:**
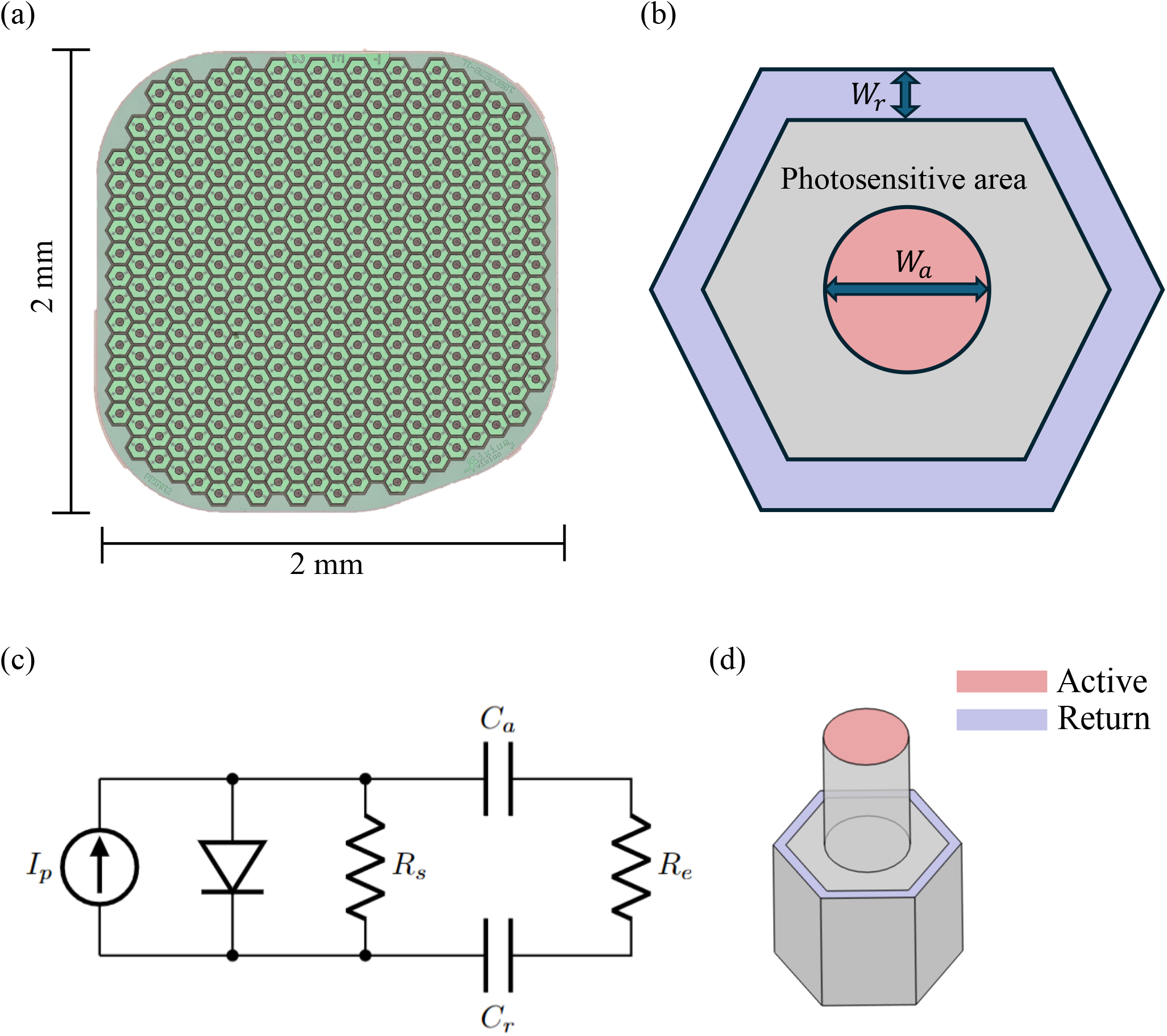
(a) PRIMA implant with 100µm wide hexagonally tiled bipolar pixels (b) Diagram of a single bipolar pixel. *W*_*r*_ and *W*_*a*_ indicate the widths of the return and active electrodes, respectively (c) The circuit model of a single pixel labelled with design parameters: *R*_*s*_, *R*_*e*_ - the shunt and electrolyte resistance; *C*_*a*_, *C*_*r*_ - capacitance of the active and return electrodes; and *I*_*p*_ - the photocurrent (d) Diagram of a bipolar pixel with a pillar electrode.

Each pixel in the PRIMA MEA is 100µm wide and contains two photodiodes connected in series, a shunt resistor, and a pair of electrodes - a central active electrode and a peripheral return electrode (Fig. 1(b,c)), a configuration that will be referred to as bipolar. The sampling limit of such an array is set by the pixel pitch, and for human retina, a 100µm spacing corresponds to a visual acuity of 20/420 [2]. Clinical results of PRIMA patients demonstrated that their prosthetic visual acuity is 20/418, closely matching this theoretical limit. With electronic zoom between the camera and the implant patients can read smaller fonts, but it comes on account of a reduced visual field [3].

Despite being the highest reported prosthetic acuity to date, further improvements in resolution are needed to help a larger population of AMD patients. Rodent models have shown that with a photovoltaic subretinal array having 40µm pixels, the grating acuity matched the pixel pitch, while with 20 µm pixels, it reached the limit of their natural resolution – 28 µm [4]. The 20 µm pixels geometrically correspond to a visual acuity of 20/80 in a human eye.

Translation of these promising results to human patients requires careful consideration of a few additional factors: 1) Unlike rodent models of retinal degeneration, human degenerate retina has a debris layer of a few tens of µm in thickness, which separates the INL from the subretinal implant [2], [5]. This layer is likely to increase the stimulation threshold and decrease both contrast and resolution, compared to rodents. 2) PRIMA’s success in eliciting form vision, as opposed to just light perception observed with many previous implants, is attributed in part to the network-mediated retinal response, which preserves many features of the natural retinal signal processing [6], [7], [8], [9]. Therefore, it is important to ensure that while the second-order neurons (bipolar cells) are stimulated at maximum strength and contrast, the third-order neurons (ganglion cells) are not activated directly.

Here, we present a multi-objective optimization of the structure and electrical parameters (e.g. pixel size, electrode configuration) of future implants that could provide higher visual acuity while maintaining selective activation of bipolar cells in human degenerate retina, and demonstrate the range of expected perceptual brightness, resolution and contrast in patients with various thicknesses of the subretinal debris layer.

## II. Methods

A pattern of NIR light projected from the AR glasses onto the photovoltaic subretinal MEA leads to the generation of electric field in tissue, which in turn drives the retinal neural response. To characterize this field, a Retinal Prosthesis Simulator (RPSim) was previously created. It combines the finite element modelling of electric fields (COMSOL) with a SPICE-based circuit solver (Xyce) to simulate non-linear circuitry of multiple pixels coupled via a common electrolyte [10], [11].

This in-house simulator is expanded in this paper as part of a multi-objective optimization pipeline with the goal of finding the optimal next-generation implant design while considering several key parameters: 1) Electrical stimulation strength, which is assessed as the voltage drop across BCs [12] and is assumed to be manifested as perceptual brightness. 2) Contrast of the electrical pattern across BCs, which should be sufficient for resolving the image. 3) Electric field in the ganglion cell layer, which should be below the threshold of direct activation of retinal ganglion cells (RGCs) and their axons. 4) The design should be robust enough to ensure proper implant performance within the expected variation in the thickness of the subretinal debris that separates the implant from the inner nuclear layer, where BCs reside. Such multi-objective optimization requires a trade-off between the various factors, and so each parameter is benchmarked to the PRIMA implant.

### A. An End-to-End Implant Evaluation Platform

Each pixel on the implant can be modeled as an electrical circuit that includes active and return electrodes, a photodiode, a shunt resistor *R*_*s*_ and a conductive medium (Fig. 1(c)). Return electrodes of all the pixels are connected, forming a continuous hexagonal mesh. At the operation frequency used in the PRIMA system (pulses of 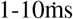 at 30 Hz frame rate) and within the voltage range of a Si photodiode (*<* 0.6 V), the dynamics of an electrode-tissue interface are closely approximated by a capacitor and resistor in series, where the capacitance of both the active electrode *C*_*a*_ and the return electrode *C*_*r*_ is determined by the sputtered iridium oxide film (SIROF) coating of the electrodes [13], [14]. The resistor *R*_*e*_ models the ohmic drop in the medium - traditionally referred to as the access resistance of the electrodes. In this diagram, the access resistances of the active and return electrodes have been combined.

Referring to the circuit model shown in Fig. 1(c), *R*_*s*_, *R*_*e*_, *C*_*r*_, *C*_*a*_, and *I*_*p*_ are all adjustable. *C*_*r*_ and *C*_*a*_ depend on the physical dimensions of their respective SIROF coatings. *R*_*e*_ depends on the geometry of the active and return electrodes. The photocurrent *I*_*p*_ depends on the photosensitive area of a pixel and light intensity, and the resistor *R*_*s*_ is adjustable in the fabrication process. The way each of these components is adjusted affects the overall performance of the implant and finding the optimal values for these variables is the essence of this paper.

The addressed electrode geometries are 1) flat, where the active and return electrodes are located on the same plane, and 2) pillars with elevated active electrodes. Flat implants with 100/µm pixels successfully provided prosthetic vision in human patients [3], while several advantages of pillar electrodes that enable smaller pixels have been modeled and demonstrated experimentally in rats [8], [15]. Here we thoroughly explore the design space of each geometry to determine the optimal configurations and the limits of their performance. This optimization process provides confidence in improving patient outcomes and is essential prior to fabricating the next-generation implant since clinical trials are so long and expensive that very few configurations can be tested.

It will be shown that *R*_*s*_ has a very shallow optimum, which does not change significantly between pillar and flat geometries. This allows treating *R*_*s*_ independently and then fixing it for examination of the other variables since its shallow optimum limits the probability that such an optimization sequence will result in a local minimum. While not ideal, this separate treatment is necessary to keep the multidimensional search space computationally tractable. To find the optimal value for *R*_*s*_, the electric field generated by Landolt C optotypes (Fig. 2(a)) was simulated using RPSim, and the stimulation strength and contrast were quantified through methods explained in section II-B below. This process was performed for arrays with 20, 40 and 55 µm pixels, and optimum values for other pixel sizes were found by interpolation.

**Fig. 2:**
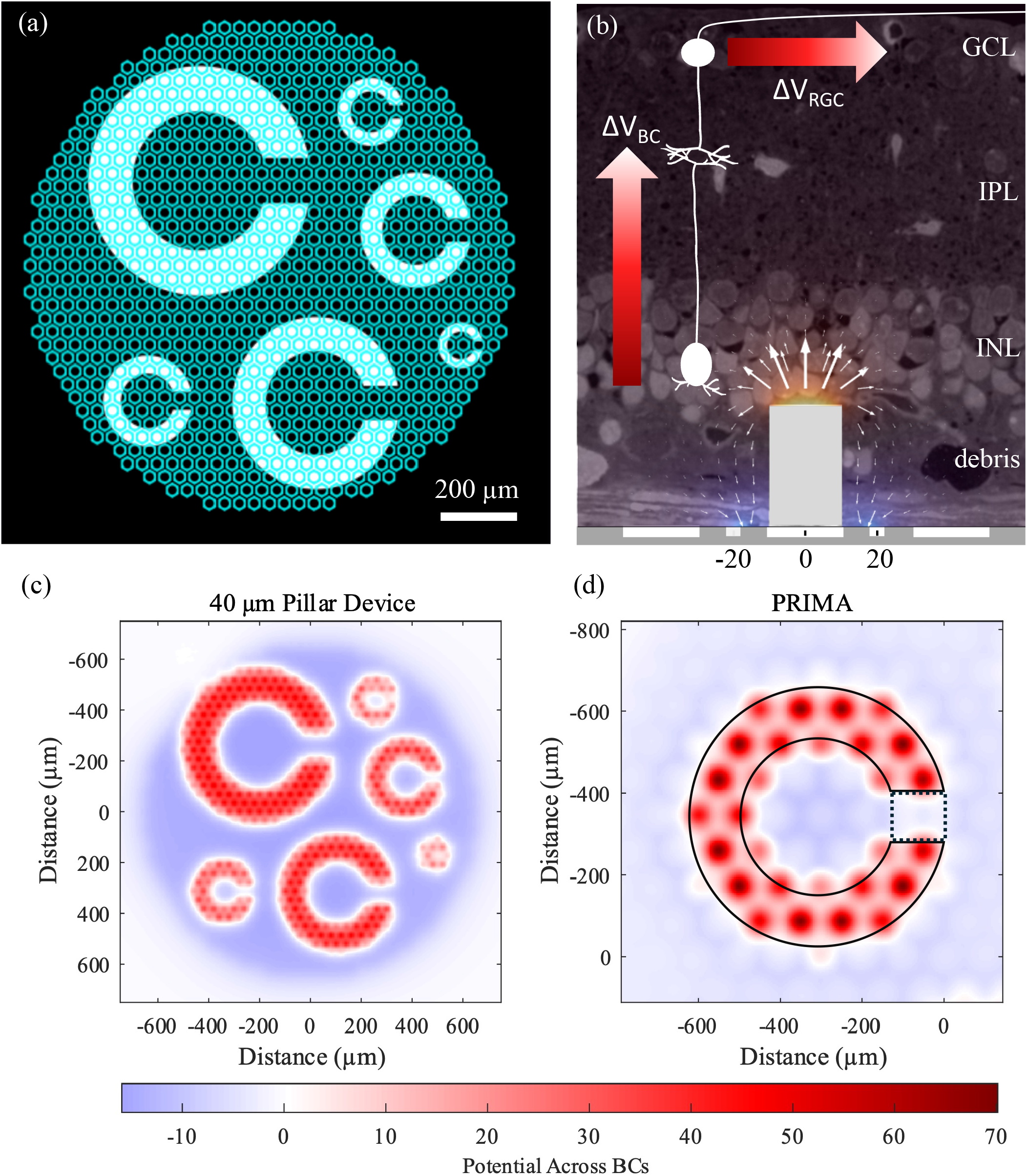
(a) Illumination pattern of tiled Landolt Cs on top of the implant containing 40µm pixels (b) Visualization of the electric potential and current density resulting from the activation of a single bipolar pixel, overlaid on histology of the retina and a diagram of a bipolar and a ganglion cell. GCL - ganglion cell layer, IPL - inner plexiform layer, INL - inner nuclear layer. Colored arrows next to the diagrammed cells indicate the planes along which the electric potential drops along bipolar cells and retinal ganglion cells (ΔV_BC_ and ΔV_RGC_). (c) ΔV_BC_ calculated from the tiled Landolt C pattern for an optimized device with 40µm bipolar pixels having pillar electrodes matching the debris layer thickness (d) ΔV_BC_ for a Landolt C with a 120 µm gap, generated by the PRIMA implant with 100µm flat pixels. The solid and dotted outlines indicate the area where the electric potential is averaged for the ‘on’ and ‘off’ portions of the pattern, used to calculate contrast.

The elementary electric fields, used as inputs to RPSim, are calculated with the finite element method platform COMSOL Multiphysics [16], and these are unique for every electrode geometry. Therefore, a COMSOL simulation must be run for every implant design under consideration. To do this, we wrote a Matlab program to automatically run a simulation with the required configuration and generate the necessary files for RPSim using COMSOL Livelink with Matlab.

### B. Evaluating Performance of a Microelectrode Array

Since PRIMA implants successfully provide form vision in human patients with an average resolution matching the pixel pitch [3], we evaluate the next-generation arrays with smaller pixels using a pattern of tiled Landolt Cs (Fig. 2(a)) – the optotypes used in visual acuity tests. To avoid the considerations of the hexagonal pixels’ orientation, sizes of these optotypes were selected such that their gap width corresponds to 1.2 times the pixel width tested. They are therefore 24, 36, 48, 60, 90 and 120 µm wide for pixels of 20, 30, 40, 50, 75 and 100µm, respectively. These patterns were projected with a pulse duration of 9.8 ms at 30 Hz frequency and irradiance of 3 mW/mm^2^, as in clinical settings [17]. The resulting electric potential in the retina (Fig. 2(c)) was then analyzed and compared with that generated by the PRIMA implant (Fig. 2(d)).

The first objective for optimization is stimulation strength, defined as the electric potential drop from bottom to top of the bipolar cells. Neuronal stimulation can be complex to model and very nuanced, however it has been shown that for retinal bipolar cells, depolarization at the axonal terminals above certain threshold activates the voltage-gated calcium channels, resulting in an increased release of glutamate into the synapses of retinal ganglion cells [12]. It has further been shown that using this metric, the stimulation thresholds measured clinically in humans closely match those recorded electrophysiologically in rats, corresponding to depolarization by about 10mV [18]. This is close to the required potential increase to open calcium channels (*>* 8mV) [19], [20], providing strong evidence that the voltage drop between the top and bottom of a bipolar cell (Fig. 2(b)) serves as a good approximation for assessment of their response and acts as a surrogate of perceptual brightness.

Despite the pixelated stimulation, patients report perceiving smooth patterns, such as lines and letters. Therefore, perceptual brightness and contrast of a pattern are likely represented by some form of average over a pattern. Due to the nonlinear nature of the cellular response to polarization, it is likely a weighted summation (e.g. the classical linear-nonlinear model) rather than just a simple integral of the potential drop between two planes over the pattern. As can be seen in Fig. 2(c), electric potential generated by the optimized arrays with smaller pixels is much more uniform than that produced by the PRIMA implant (Fig. 2(d)). Since potential within the outlined pattern in Fig. 2(d) is above the stimulation threshold, and the maximum-to-average ratio is significantly higher with the PRIMA implant, the maximum rather than average over the pattern serves as a worst-case estimate for new implants, when compared to PRIMA. To avoid complex assumptions regarding the nonlinearity of cellular response and to provide a conservative comparison of the new implants to PRIMA, we assess the pattern brightness by the maximum potential drop across bipolar cells within it, which are located above the active electrodes.

The second metric is the contrast of the resulting electric field pattern, as ‘sensed’ by the BCs. To resolve the gap between the projected lines of a letter, the electric field must rapidly decline near the edges of the letter. The contrast *C* is defined as the relative difference between the average stimulation strength of the Landolt C pattern *I*_*p*_ and its gap *I*_*g*_ (Fig. 2(d)): *C* = (*Ip* −*Ig*)*/*(*Ip* + *Ig*).

The third metric is neural selectivity. We aim to selectively stimulate BCs while avoiding the direct activation of RGCs, and especially of their axons, which would distort the retinotopic map of the percepts [21], [22]. We discuss the best predictor for axonal stimulation further in II-C. Since PRIMA patients did not report distortions in prosthetic vision, we assume that they do not experience axonal stimulation. With new implants, we first optimized towards high neural selectivity, and then scaled the irradiance such that the drop in electric potential in the nerve fiber layer (NFL) never exceeds 20 mV, roughly matching that from PRIMA.

Fourth, optical coherence tomography (OCT) imaging demonstrated that, unlike rodents, in human patients the INL is separated from the implant by a layer of debris [2]. Its thickness varies between patients and within individual retinas in the range of about 25 – 55 µm [5]. To assess the effect of the debris layer on stimulation strength and attainable resolution, we shift the location of the retinal cell by the debris thickness. For flat implants, we used a range of 20 to 57 µm. For pillar implants, their height was designed to match the average debris thickness of 35 µm, and we varied the debris thickness above the pillars from an additional 2 to 22 µm (corresponding to the total of 57 µm). The BCs are modelled as 52 µm tall cells, starting above the debris, and the NFL is located 31 µm above the top of the BCs.

Finally, it is important to note that since electric potential is affected by crosstalk of neighboring pixels, these metrics depend on the size of the projected image. A Landolt C is considered resolvable if the stimulation strength exceeds the voltage corresponding to the highest clinical thresholds (16.4 mV [10]), and the contrast exceeds that with the PRIMA implant, calculated with debris thickness of 37 µm.

### C. Estimating the Threshold of Axonal Stimulation

The full-fledged modeling of a neuron’s response to an extracellular electrical stimulus (as computed using Neuron [23]), requires a detailed map of the cell shape and its ion channels. To provide a more universal estimate, several approximations have been proposed over the last decades, based on characteristics of the electric field rather than a particular cell configuration.

The well-known “activating function”, developed by Rattay [24], suggests that axonal stimulation is proportional to the second derivative of electric potential along the axon. However, subsequent literature raised several limitations of this estimate. A more recent study showed that for axon length in a mm range, stimulus duration in ms range, and distance from electrode of hundreds of µm, a better predictor might be the inverse of the external electric potential centered at the mean – called the “mirror” estimate [25].

For subretinal stimulation of human RGCs in the macula, the distance from the implant to NFL is about 120 µm [5], and our stimuli last 1-10ms. Therefore, in some scenarios the “mirror” estimate may be more appropriate than “activating function” for assessing axonal stimulation in RGCs. However, this approximation was only validated with a point source and a uniform electric field. In subretinal stimulation with photovoltaic arrays, the electric field is far more complex since it replicates an image projected onto the retina and may include several peaks and valleys along an axon.

A third metric for predicting neural response is the voltage drop along the cell, which we will call the “voltmeter” approximation, calculated as the difference in electric potential between its dendritic and axonal ends. This metric was shown to be accurate for retinal bipolar cells. [12].

To determine the best metric for predicting axonal stimulation of RGCs in a complex electric field, like the one generated by a multielectrode array, we compared the threshold calculated by a full-fledged Neuron model with the three aforementioned approximations. Axonal stimulation depends heavily on the spatial variation of electric field relative to the axon. To account for this, four different stimulation patterns were evaluated, and axons were placed to cross the areas with the largest variation in electric potential, as shown in Fig. 3. A Landolt C with a 120 µm stroke width, a 400 µm wide bar, a 1000 µm wide rectangle covering the top half of implant, and a 1600 µm diameter circle.

**Fig. 3:**
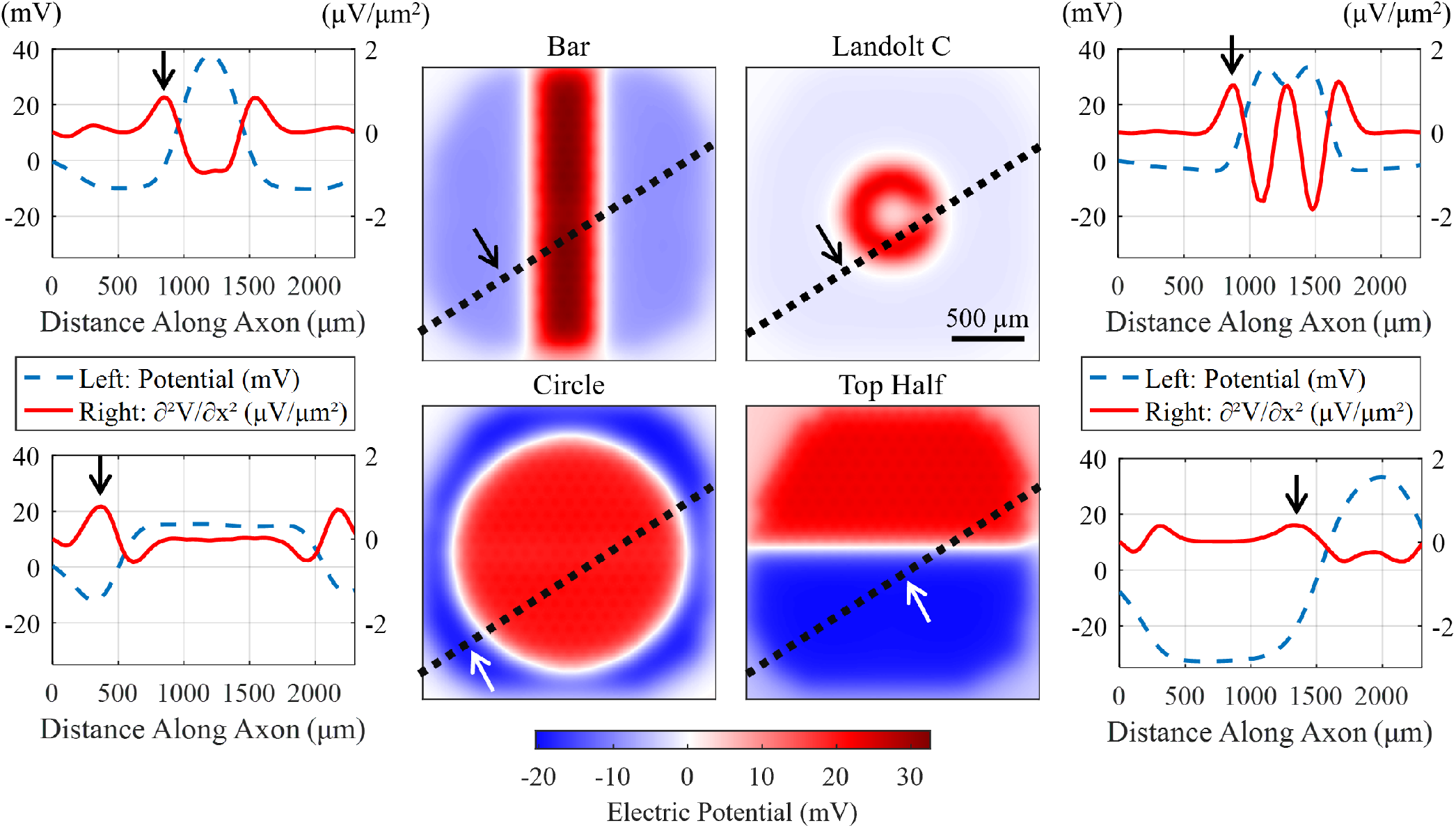
Electric potential in the nerve fiber layer (120 µm above the implant) generated by the 2×2mm PRIMA implant used in human patients. Patterns are generated using an irradiance of 3 mW/mm^2^ and the electric potential is scaled linearly to find the stimulation threshold. Dotted lines over the patterns show the position of the retinal ganglion cell axon. The extracellular electric potential (dash, blue) and its second derivative along the axon (solid, red) at threshold are plotted next to each pattern. Arrows indicate the spike initiation site observed in Neuron.

We placed a human midget RGC, the most common RGC in the macula [9], adjusted to have a 10 mm long axon 120 µm above the electrode array. To explore only the axonal stimulation thresholds, we placed both ends of the cell (the cell soma with its axonal initial segment, and the end of the axon) outside of the strong electric field. To mimic the typical stimuli with photovoltaic implants [17], we applied 10 ms long monophasic pulses at 30 Hz repetition rate. Modeling was performed in the Neuron 7.2 software, with a time integration step of 0.01 ms. The stimulation started after 50 ms and lasted for 2 seconds. The threshold was defined as the pulse amplitude resulting in a 50% spiking probability, found by fitting a sigmoid using MATLAB’s 4 parameter logistic model.

At the threshold, we used the external electric potential along the axon to calculate the following: 1) The second derivative of electric potential *∂*^2^V*/∂*x^2^ along the axon; 2) The “mirror” approximation V_mean_ − V_min_; and 3) The “voltmeter” approximation V_max_ − V_min_. As seen in Fig. 3, the spike is initiated near the maximum of the second derivative of electric potential, as predicted by the “activating function”, and near the minimum of electric potential as predicted by the “mirror” estimate and “voltmeter” approximation. However, there was a large spread of stimulation thresholds between various patterns, listed in Table I. To determine which one had the least variation among various patterns and therefore provides the most accurate prediction for axonal stimulation in general, we compared the maximum and minimum values with various patterns for each metric. It was found that the “voltmeter” approximation performed the best, with a ratio of 2.4 between max and min, whereas the second derivative and “mirror” approximations differed by factors of 4.3 and 34.2, respectively.

**TABLE I:** Three approximations of the axonal stimulation threshold of retinal ganglion cells with spatially varying stimuli. The maximum of the second derivative of electric potential, “mirror”, and “voltmeter” estimates at the threshold for axonal stimulation found with NEURON. The variance indicates how accurate a given metric is with various stimulation patterns.

| Estimate | Bar | Circle | Landolt C | Top Half | Variance (max/min) |
| --- | --- | --- | --- | --- | --- |
| **Max** $\partial^2 V / \partial x^2$ [$\mu V / \mu m^2$] | 1.0 | 0.9 | 1.7 | 0.4 | 4.3 |
| **Vmean-Vmin** (mirror) [mV] | 9.0 | 8.7 | 1.1 | 37.6 | 34.2 |
| **Vmax-Vmin** (voltmeter) [mV] | 48.1 | 27.7 | 42.5 | 65.67 | 2.4 |

While the “voltmeter” approximation performs the best among the three approximations, a factor of 2.4 is still a large variation, suggesting that simple numerical metrics regarding the electric field may not be able to accurately predict subretinal axonal stimulation of RGCs with complex electric fields. However, since this large variation is primarily due to the spatial characteristics of the stimulating pattern, by eliminating this variable, we can safely use the “voltmeter” approximation as the metric for the implant optimization. I.e. the implants are optimized for minimizing the axonal stimulation using the same pattern for comparison. Since PRIMA implant provides form vision in patients, we assume that its electric field does not cause axonal stimulation, which would result in distorted percepts [22]. Therefore, we can use the PRIMA implant as the benchmark of the acceptable electric field for comparison with the new devices.

Using several stimulation patterns, we calculated the potential difference along the axon, i.e. the “voltmeter” approximation, from the PRIMA implant and an optimized implant having 40µm wide pixels with pillars at 3 mW/mm^2^. We then selected the worst pattern, i.e. the one with the largest ratio of electric potentials (ΔV_new_*/*ΔV_Prima_). To avoid axonal stimulation, we then reduced the stimulus strength (irradiance) such that ΔV_new_ = ΔV_Prima_. Since for all other patterns ΔV_new_*/*ΔV_Prima_ is lower, they avoid axonal stimulation. As shown in Table II, among the four patterns, the ratios range from 1.44 to 1.91, with the Landolt C pattern having the largest. Therefore, we used the “voltmeter” approximation with a Landolt C having a 120 µm stroke width for optimization of the neural selectivity of new photovoltaic arrays. Clinical trials used irradiances up to 3.5 mW/mm^2^, which yields a 17 mV drop along RGC axons. Since this is well below the “voltmeter” approximation thresholds from Table 1, we rounded to 20 mV as a rough estimate of the maximum value allowed in further assessments of the implants’ performance.

**TABLE II:** Maximum possible voltage drop along an axon in any orientation calculated using the “voltmeter” approximation for several different illumination patterns. The corresponding electric fields were generated by the PRIMA implant (ΔV_Prima_), and by an optimized device with 40µm pixels and pillar electrodes (ΔV_new_) at an irradiance of 3 mW/mm^2^. The ratio is the voltage drop generated by the optimized device to that of the PRIMA implant.

| Pattern | $\Delta V_{\text{Prima}}$ [mV] | $\Delta V_{\text{new}}$ [mV] | Ratio [V/V] |
| --- | --- | --- | --- |
| **Bar** | 26.3 | 45.5 | 1.73 |
| **Circle** | 26.6 | 38.2 | 1.44 |
| **Landolt C** | 14.6 | 28 | 1.91 |
| **Top Half** | 30.2 | 48.7 | 1.61 |

### D. Monopolar Pixels

One method for reducing the pixel size is to employ a monopolar (MP) electrode design. This involves replacing the local return mesh in each pixel with one global electrode exposed to the medium at the periphery of an implant (Fig. 4) [10]. With this approach, the uniform activation of a disk pattern results in an electric field similar to that of a correspondingly sized disc electrode. Summation, or crosstalk, of electric fields from neighboring monopolar pixels allows generating stronger electric fields than bipolar pixels, where local electric fields are much more confined. However, much stronger crosstalk between neighboring pixels also leads to significantly lower contrast and higher electric field in the NFL.

**Fig. 4:**
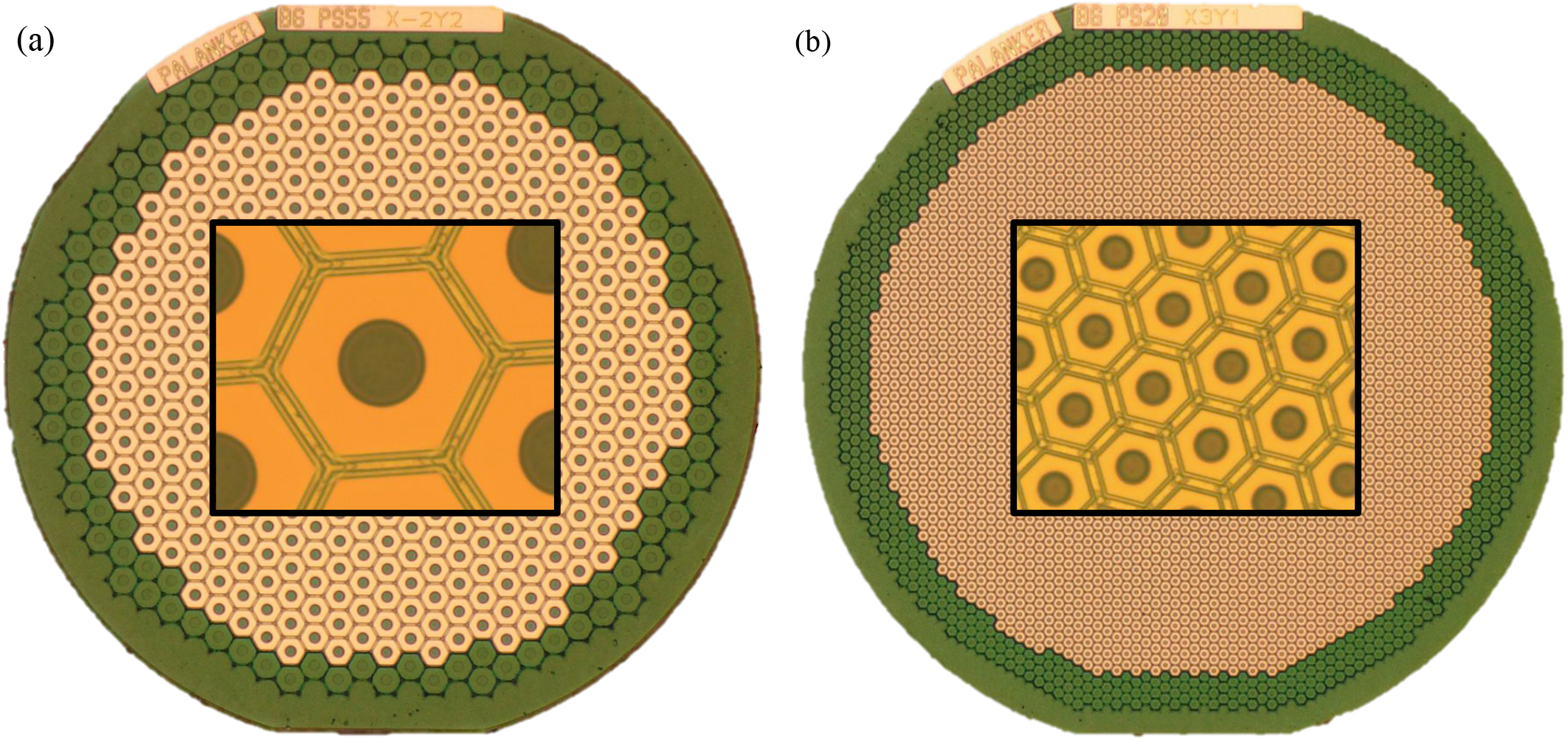
Monopolar arrays of 1.5 mm in diameter, composed of 55 (a) and 20 µm (b) pixels. In the central area of the arrays, connected return electrodes are hidden under the insulation layer. The green ring along the edge of the implant is the exposed common return electrode. Insert in the middle of each image shows a higher magnification view of the pixels.

Lowering the capacitance of the global return electrode forces the electric current to return through the shunt resistors of the non-illuminated pixels and through the diodes of the pixels charged during the previous pulse. This results in a more constrained electric field, as current does not flow as far on a return path, leading to better contrast and selectivity. Assuming a fixed capacitance of SIROF coating on active electrodes, we can perform a one-dimensional sweep of the return electrode capacitance to examine the effects of forcing more of the current to return through the nearby dark pixels. It was shown earlier that for the same pattern, arrays of 20 µm pixels provide higher contrast, but lower stimulation strength compared to larger pixels [10]. Since the main drawback to the monopolar design is contrast, in this optimization, we sampled the return-to-active capacitance ratio from 0.01 to 5 for implants with 20 µm pixels.

### E. Bipolar pixels

For bipolar pixels, *R*_*a*_, *C*_*r*_, *C*_*a*_, and *I*_*d*_ can be optimized with a two-dimensional parameterized sweep, where parameters are the relative areas of the active and return electrodes in the pixel. This fully defines a pixel’s geometry based on the radius of the active electrode, *W*_*a*_, and the width of the return, *W*_*r*_ (Fig. 1(b)). Since the minimal width of the return electrode *W*_*r*_ is limited to 2 µm by design rules in the fabrication processes, smaller return capacitances are simulated using a thinner SIROF layer (lower specific capacitance). Inability to reduce the return electrode width below 2 µm harms performance of small pixels (starting at 30 µm for flat implants and 50 µm for pillars) as it decreases the photosensitive area and thereby the photocurrent. If the resolution of the fabrication process will improve, photoresponsivity of small pixels should increase and match that of the larger pixels. As a result, smaller pixels would perform a little better than is predicted here. The optimization presented can also be used directly for the optimization of MEAs utilizing emerging technologies and materials such as 3D flexible electrodes [26], and coatings such as upconversion nanoparticles which may improve the photoelectric conversion efficiency [27], [28], improving performance even further.

In search of the optimum, we varied the active and return electrode sizes from 5 to 40% of the pixel area on 20, 30, 40, 50, and 75 µm pixels. These sizes were chosen to correspond to devices that have already been fabricated, and they provide a dense enough sampling for assessing the best possible resolution. From the results of this grid search, a Pareto frontier was formed and an optimal implant was selected for each geometry with respect to the four key design parameters.

## III. Results

### A. Shunt resistance

The first optimized parameter was the value of the shunt resistor, *R*_*s*_. Since it provides a path of return for current via neighboring dark pixels, it helps constrain the electric field and improve the image contrast. Shunt resistors also help discharge the electrode capacitance between the pulses and thereby increase the injected current during the next pulse, thus increasing the stimulation strength [29]. On the other hand, excessive drainage of the photocurrent through the shunt of the same pixel limits the current flowing into the tissue. Balance of these opposing effects results in an optimal shunt value when considering the stimulation strength (Fig. 5(a)).

**Fig. 5:**
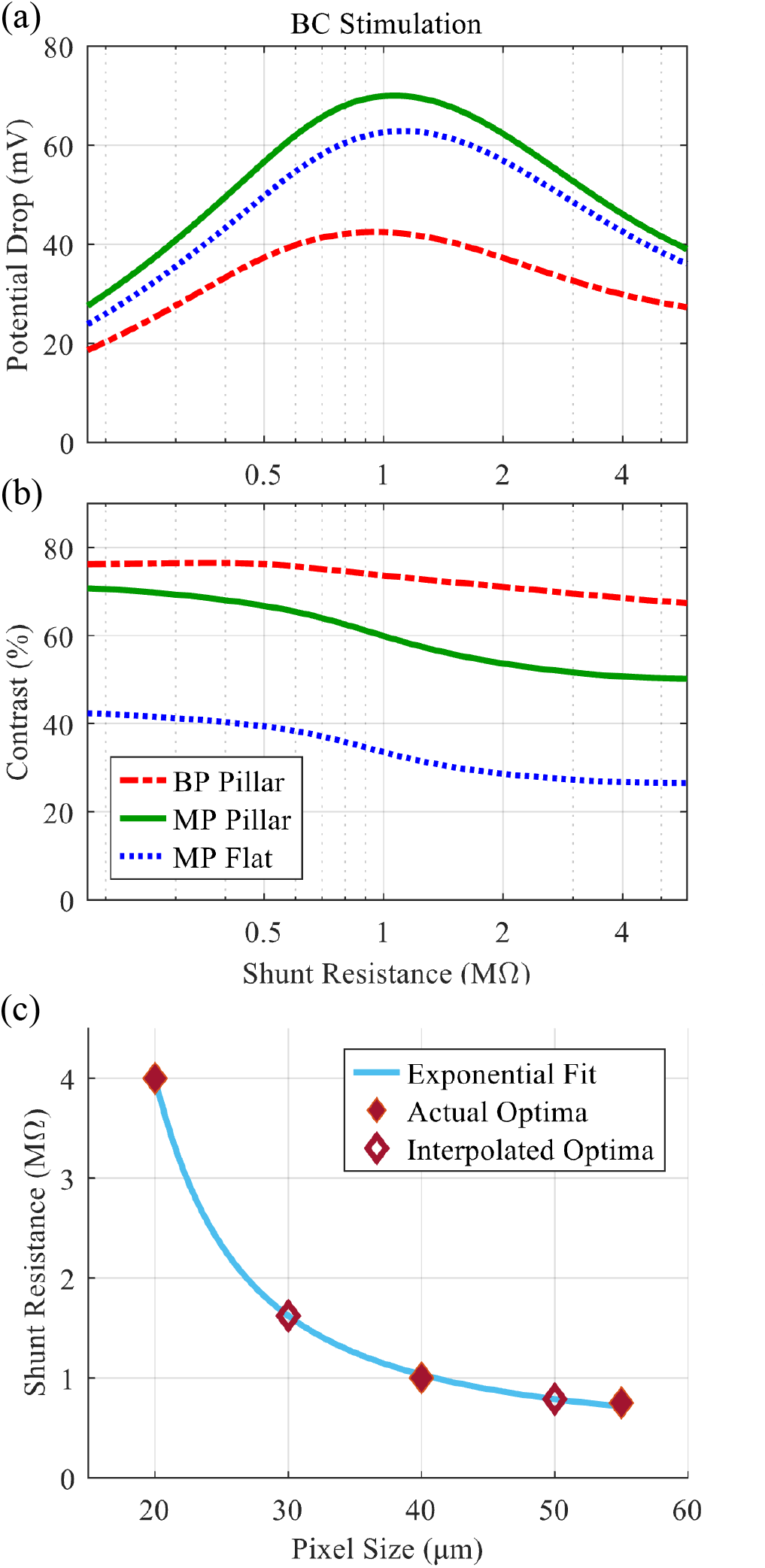
(a) The drop in electric potential across bipolar cells (BC) and (b) contrast as a function of the shunt resistance, calculated for bipolar (BP) pillar electrodes and monopolar (MP) pillar and flat electrodes, all with 40µm pixels (c) Optimal shunt resistance that provides the highest stimulation strength for various pixel sizes.

We determined optimal *R*_*s*_ values for implants with monopolar and bipolar electrode configurations, of flat and pillar geometries. For 40µm pixels, the optimal shunt value for maximizing the stimulation strength varies at most by 11% from the average of all 3 scenarios, which was around 1 MΩ (Fig. 5(a)). Due to the extremely shallow shape of this optimum, using this average value results in less than 0.4% drop of the stimulation strength compared to the exact optima for each scenario, meaning we can safely use one shunt resistance for a pixel size regardless of the specific electrode geometries. We also found no significant changes in this value nor performance loss due to variation in retinal resistivity from 200 to 3000 Ω * cm, nor for different projection patterns. The contrast for a Landolt C with a 48 µm gap is shown in Fig. 5(b). Lower shunt resistance provides higher contrast (due to better field confinement), but weaker stimulation. For example, lowering the shunt resistance to 0.5 MΩ improves the contrast of MP pillar devices from 60 to 67%, but results in a 20% decrease in stimulation strength. For the other electrode geometries, the tradeoff is even worse.

A similar process was performed for 20 and 55 µm pixels, resulting in the optima around 4 and 0.75 MΩ, respectively. Shunt resistances for 30 and 50 µm pixels were then found by interpolation (Fig. 5(c)).

### B. Monopolar arrays

In the monopolar array, the local returns in each pixel are hidden under the insulator and a common return electrode covers the circumference of the implant (Fig. 4). Summation of electric fields from multiple illuminated pixels increases the penetration depth of electric field into the tissue and reduces the contrast of the pattern, which can lead to poor neural selectivity and strong variation of the stimulation strength with the size of the projected image. As was discussed in section II-D, the stimulation selectivity and contrast can be improved by minimizing the return capacitance. This forces the current to return through local shunt resistors of inactive (dark) pixels, at the cost of lower stimulation strength. Since contrast is the limiting factor for this design, we maximized it by using the smallest modeled ratio of the return/active capacitance: 0.01. This effect largely plateaus at a ratio below 0.04, at which point any additional decrease in this ratio results in a very marginal change of electric field. However, practically removing the global return in this manner comes with additional challenges as it limits the maximum illuminated area on the device (see details in Supplementary Information).

The expected performance for the 20 µm monopolar array with both flat and pillar electrodes is shown in Fig. 6. Stimulation strength and pattern contrast for BCs, as well as the potential drop in the RGC layer, are plotted as a function of the Landolt C gap size, calculated for 3 debris thicknesses. Irradiance was scaled such that the electric potential drop in RGC layer never exceeds 20 mV (see Tables III and IV). The performance of the PRIMA device in the best case (20 µm debris) and worst case (37 µm debris) is shown by horizontal dash lines for comparison, although it applies only to the largest font size (120 µm gap). Finally, the corresponding stimulation thresholds of bipolar cells (best and worst, based on clinical data [10]) are shown by colored strips. Because the irradiance must be scaled down significantly with pillar electrodes to avoid RGC stimulation, they do not offer a significant improvement in BC stimulation strength compared to their flat counterparts. They do, however, provide higher contrast. The stimulation of RGCs is nearly 4 times weaker with the smallest pattern than the largest. This implies that if large patterns could be avoided, the implant irradiance could be increased, thus providing significantly stronger stimulation of bipolar cells. This could be achieved by creating sparse visualizations of objects. Similarly, this reduction in irradiance could be foregone altogether if it is found that the stimulation of RGCs does not interfere with the retinotopic map.

**Fig. 6:**
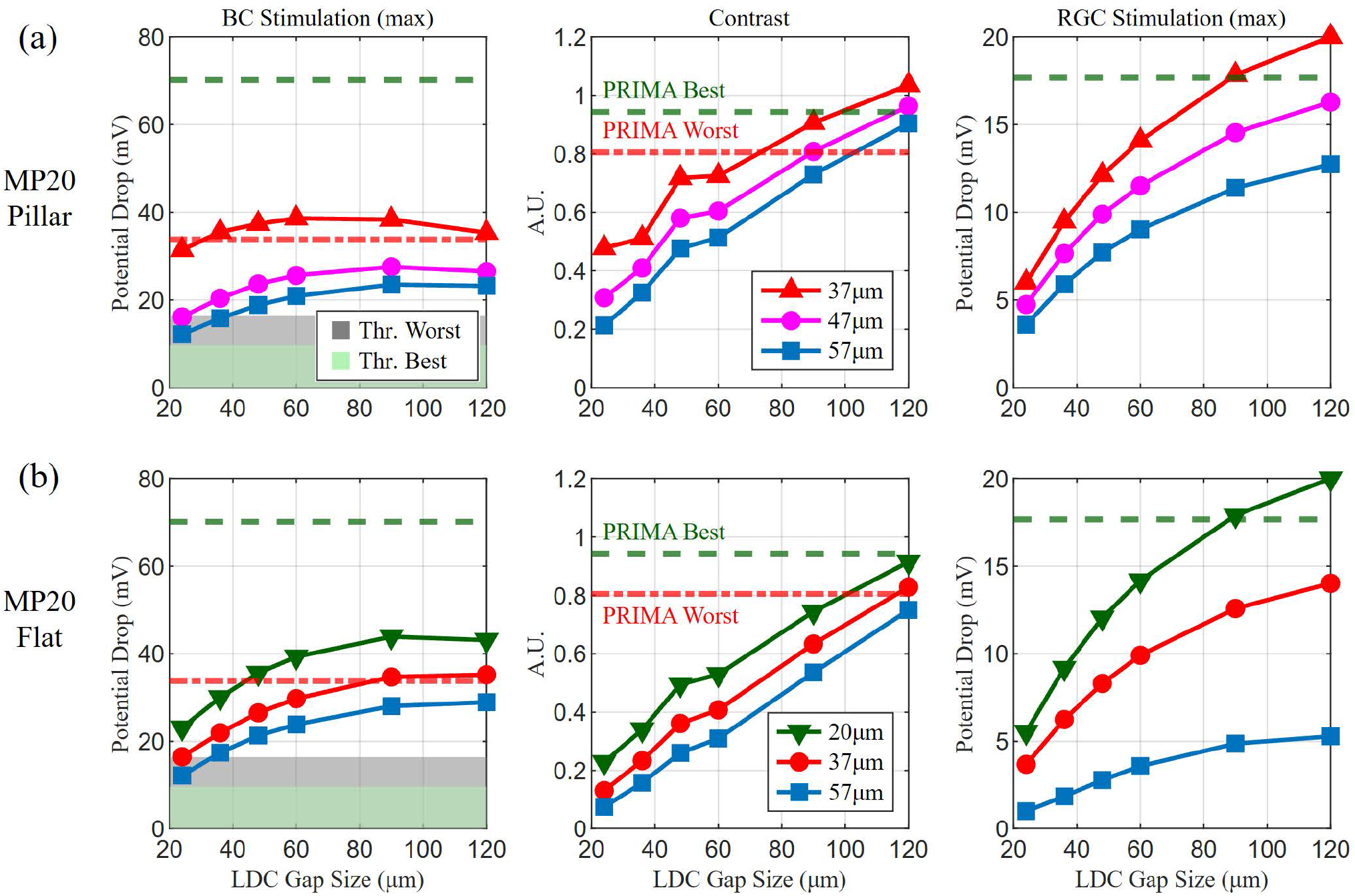
Performance of monopolar (MP) arrays for the maximum potential drop over a pattern.(a) Bipolar cell (BC) stimulation strength, contrast and potential drop in the retinal ganglion cell (RGC) layer as a function of the letter size (gap width in Landolt C) projected onto monopolar array with 20 µm pixels having pillar electrodes of 35 µm in height. Red, magenta and blue lines correspond to debris thickness of 37, 47 and 57 µm, respectively. Green and grey shaded zones show the stimulation thresholds of bipolar cells at 10 ms for the best and worst patients (b) Similar plots for flat arrays. The additional green plot corresponds to 20 µm debris thickness.

**TABLE III:** The predicted minimal resolvable Landolt C gap sizes, in µm, for each evaluated implant design with flat electrodes and the laser irradiance used for each device to maintain the retinal ganglion cell voltage drop below 20 mV. Italic font indicates values of resolution worse than 120 µm, and excluded from the average value. Bold font highlights the best possible performance for a given debris thickness.

|  | Monopolar flat |  | Bipolar flat |  |  |  |  |
| --- | --- | --- | --- | --- | --- | --- | --- |
| Pixel size ($\mu\text{m}$) | 20 | 40 | 20 | 30 | 40 | 50 | 75 |
| Irradiance ($\text{mW}/\text{mm}^2$) | 2.5 | 2.0 | 3.0 | 2.6 | 2.2 | 2.2 | 2.3 |
| 20 $\mu\text{m}$ debris | 101 | 113 | 41 | **38** | 48 | 60 | 90 |
| 37 $\mu\text{m}$ debris | 116 | *120+* | 73 | 67 | 66 | **62** | 90 |
| 47 $\mu\text{m}$ debris | *120+* | *120+* | 91 | **77** | 83 | 89 | 90 |
| 57 $\mu\text{m}$ debris | *120+* | *120+* | 112 | **95** | 103 | 106 | 112 |
| Average | 109 | 113 | 79 | **69** | 75 | 79 | 96 |

**TABLE IV:** The predicted minimal resolvable gap sizes, in µm, for each implant design with pillar electrodes of 35 µm in height and the laser irradiance used for each device to maintain the retinal ganglion cell (RGC) voltage drop below 20 mV. Bold font indicates the best possible performance for a given debris thickness.

| Pixel size ($\mu\text{m}$) | Monopolar Pillar | | Bipolar Pillar | | | | |
| --- | --- | --- | --- | --- | --- | --- | --- |
|  | 20 | 40 | 20 | 30 | 40 | 50 | 75 |
| Irradiance ($\text{mW}/\text{mm}^2$) | 1.5 | 1.3 | 2.2 | 2.2 | 2.1 | 2.0 | 2.0 |
| 37 $\mu\text{m}$ debris | 73 | 96 | **24** | 36 | 48 | 60 | 90 |
| 47 $\mu\text{m}$ debris | 89 | 106 | 48 | **36** | 48 | 60 | 90 |
| 57 $\mu\text{m}$ debris | 103 | 115 | 56 | 55 | **54** | 60 | 90 |
| Average | 88 | 106 | 43 | **42** | 50 | 60 | 90 |

While the stimulation strength remains above the threshold with smaller Landolt Cs, the contrast falls below the PRIMA benchmark, which limits the utility of these devices. In the flat configuration, a 100µm Landolt C can be resolved for patients with a thin (20 µm) debris. By employing pillar electrodes, the minimum size of a resolvable Landolt C with debris matching the pillar height is 73 µm – a 1.62 times improvement in resolution from the PRIMA implant. Even optimized, monopolar implants are still unable to provide sufficient contrast and neural selectivity to offer dramatic improvements in resolution (see details in Tables III and IV).

### C. Bipolar Arrays

Bipolar arrays have similar shape and size to monopolar devices, but as with the PRIMA implant shown in Fig. 1(a), a mesh of local returns in each pixel is exposed to electrolyte and there is no ring of a common return along the edge of the implant. For bipolar implants, we first found the optimum sizes of the active and return electrodes in flat arrays and those with pillar electrodes. This was accomplished with a uniform grid search. Suboptimal designs were eliminated using Pareto optimality with the objectives of maximizing the stimulation strength and the neural selectivity. This process is described in detail in Supplementary Information. For most flat implants, the best performance occurs when allocating 25% and 12.5% of the pixel area to the active and return electrodes, respectively. The only exception was for 20 µm pixels, which performed better with 30% and 10% respective allocations. Pillar implants all performed optimally with a 25% active and 20% return area allocation. For each design, the irradiance was scaled such that electric potential in the NFL remained below the axonal stimulation threshold (see details in Tables III and IV).

Fig. 7(a,c,d,f) depicts the stimulation strength and contrast calculated for the Landolt Cs corresponding to 1.2 times the pixel width. In Fig. 7(b,e) the stimulation strength is plotted for a Landolt C of a 120 µm gap – the size resolvable with the PRIMA implant. The rightmost point on each plot shows the performance of the PRIMA implant, indicated by an empty symbol as it is not a newly optimized device. The debris thicknesses and thresholds are indicated in the same manner as in Fig. 6. On the plots for pillar devices, a horizontal green dash line indicates the PRIMA best case (20 µm debris) performance. With both flat and pillar implants, the stimulation strength diminishes quickly with decreasing pattern size. However, pillar implants maintain stimulation above the threshold for much smaller patterns than their flat counterparts. Additionally, performance with a 120 µm Landolt C shows that percepts provided by pillar implants with small pixels will not be any dimmer than with the PRIMA device for any patterns the PRIMA implant can resolve at any debris thickness. Contrast with both flat and pillar devices also exceeds that of PRIMA for all patterns which stimulate above the threshold. This means that stimulation strength rather than contrast limits the resolution of flat bipolar implants.

**Fig. 7:**
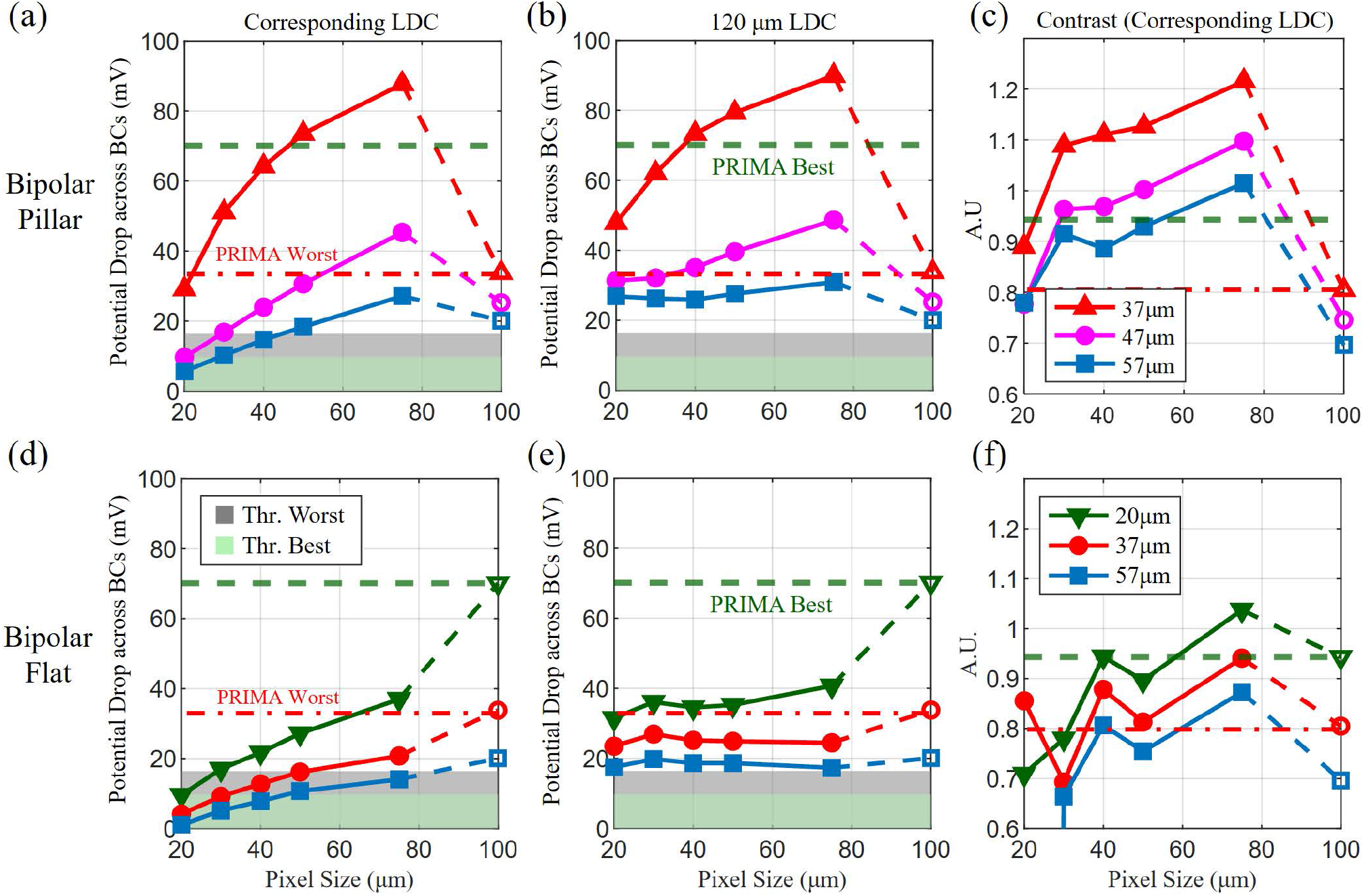
Performance of bipolar arrays for the maximum potential over a pattern. (a) Bipolar Cell (BC) stimulation strength for bipolar arrays with 35 µm tall pillars as a function of pixel size when Landolt C (LDC) gap width corresponds to 1.2 times the pixel width. (b) BC stimulation strength as a function of pixel size when Landolt C size is fixed to 120 µm (c) Contrast of the pattern in BC layer corresponding to conditions in (a). Red, magenta and blue lines correspond to debris thickness of 37, 47 and 57 µm, respectively. Green and grey shaded zones show the stimulation thresholds of bipolar cells at 10 ms for the best and worst patients. Horizontal green dash line labeled “PRIMA best” indicates the performance of the PRIMA device with 20 µm debris (d, e, f) Same as in (a, b, c), but for flat bipolar pixels, with the green line showing debris thickness of 20 µm. The rightmost points on all plots indicate the performance of the PRIMA device.

Resolution limits depend on the implant geometry as well as the thickness of the debris layer and are summarized in Tables III and IV. For flat implants and 37 µm thick debris, the smallest resolvable Landolt C is 62 µm - achieved by a device with 50 µm pixels. Since the stimulation strength grows dramatically with proximity to the target neurons, pillar electrodes allow better performance in nearly all respects. With a pillar height matching the debris thickness, the achievable resolution matches the sampling limit of the pixel pitch down to 20 µm. Smaller pixels therefore have the advantage when it comes to the best-case anatomies. However, as the debris thickness exceeds the pillar height, resolution declines faster with smaller pixels. For a 57 µm debris layer, 40µm pixels provide the best possible resolution of a 54 µm Landolt C gap. When considering all three debris thicknesses, 30 µm pixels provide the smallest resolvable Landolt C with an average gap width of 42 µm. If the debris layer thickness varies across the implant, patients may have slightly better or worse visual acuity in different parts of their visual field.

As mentioned in Section II-B, we used the maximum drop in electric potential across bipolar cells as a conservative estimate to quantify perceptual brightness of an image. Despite the pixelated stimulation, patients report perceiving smooth patterns, such as lines and letters. Therefore, perceptual brightness and contrast of a pattern might be represented by some form of average over a pixel or a pattern. Similarly, there is some evidence that stimulation thresholds of the visually evoked potentials in rats correspond to the population average over a pixel rather than the maximum of electric potential above an electrode [30]. Using the potential drop across bipolar cells, averaged over a pixel or over a pattern, as a measure of the stimulation strength or perceptual brightness significantly changes the expected performance of the implants. As shown in Fig. 8(d) for flat bipolar arrays, the average stimulation strength with smaller pixels is similar to or exceeds that of PRIMA implants for 120 µm Landolt C, unlike a sharp decline with the pixel size shown in Fig. 7(e) for the maximum values. Also, with smaller patterns (corresponding LDC, Fig. 8(c)), the decline with the pixel and letter size is much slower than in Fig. 7(d). Similarly, performance of pillar implants with larger patterns (120 µm LDC, Fig. 8(b)) is less dependent on the pixel size compared to Fig. 7(b), and with smaller letters (Fig. 8(a)) the decrease with pixel size begins only below 40µm.

**Fig. 8:**
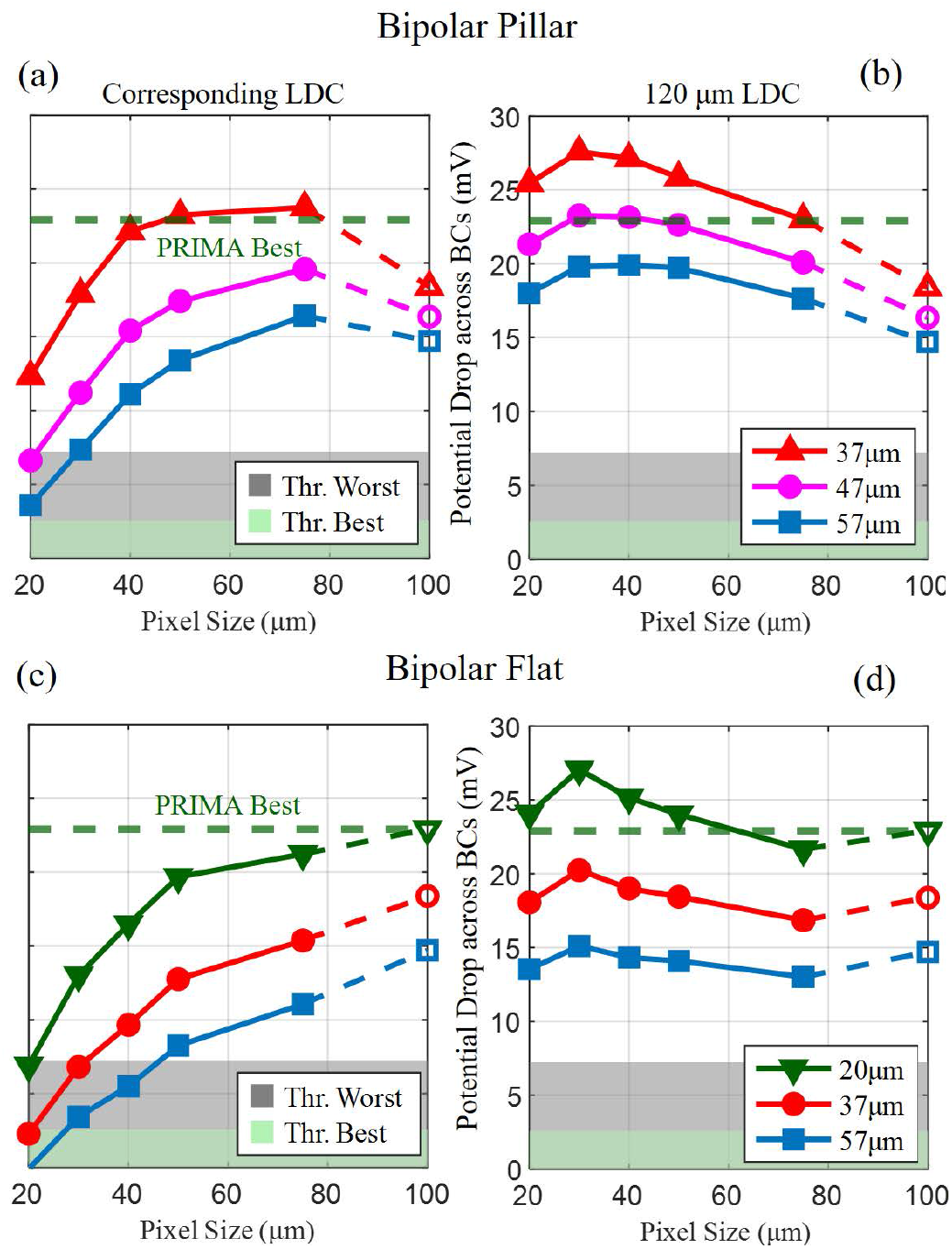
Performance of bipolar arrays with the potential drop averaged over the stimulating pattern. (a) Bipolar cell (BC) stimulation strength, for bipolar arrays with pillar electrodes as a function of pixel size when Landolt C (LDC) gap width corresponds to 1.2 times the pixel width (b) same as (a) but with the Landolt C gap size fixed to 120 µm. Red, magenta and blue lines correspond to debris thickness of 37, 47 and 57 µm, respectively. Green and grey shaded zones show the stimulation thresholds, averaged over the stimulation pattern, of bipolar cells at 10 ms for the best and worst patients. Horizontal green dash line labeled “PRIMA best” indicates the performance of the PRIMA device with 20 µm debris. The rightmost points on all plots indicate the performance of the flat PRIMA device (c, d) The same plots for flat bipolar arrays. Green, red and blue lines correspond to debris thickness of 20, 37 and 57 µm, respectively.

## IV. Discussion

Previously proposed configurations of subretinal arrays have offered dramatic improvements, but do not capture all of the necessary aspects for high fidelity prosthetic vision in patients impaired by geographic atrophy. Monopolar arrays, which achieved resolution matching natural in rodents [4], [10], are likely to provide poor contrast and low brightness in humans due to the debris layer (Fig. 6(b)). Honeycomb electrodes offer exceptional stimulation strength and neural selectivity in rodents [31], but may be filled with the subretinal debris instead of the intended INL in human patients, which would limit their efficacy. Pillar electrode arrays also demonstrated resolution matching their pixel pitch in rodents [8], [15]. To provide realistic clinical expectations, in this study, we assessed the effect of the debris layer separating the implant from INL (20-57 µm) seen in actual human patients [2], [5], as opposed to an animal model lacking this layer. We also considered neural selectivity, which was not taken into account previously. The present study addresses these factors as a function of the pixel size, aiming to maximize stimulation strength and contrast without stimulating the RGC axons.

Optimization of the return electrode size in bipolar implants can be viewed as a gradual transition from the highly constrained and localized electric field of bipolar pixels with large return electrodes, such as in PRIMA, to the strong and deep-penetrating field of a monopolar array with no local return electrodes. We demonstrated earlier that scaling the PRIMA pixels down is challenging: to provide the same stimulation strength with 75 µm pixels as that with 100µm pixels of PRIMA, the laser’s irradiance should increase from 3 to 8 mW*/*mm^2^ - approaching the thermal retinal safety limit, which is 8.25 mW*/*mm^2^ for an ocular device of this size and a 30% duty cycle [18]. Relaxing the electric field confinement by decreasing the return electrode capacitance increases the stimulation strength but decreases the contrast and selectivity of neural stimulation. In this study, we demonstrated the expected performance in optimal configurations of subretinal implants with pixel sizes down to 20 µm. Due to strong crosstalk between neighboring pixels in monopolar arrays, performance of such implants is significantly worse than that of bipolar arrays. However, there are additional controls which may improve their characteristics. One of them is a method of contrast enhancement by ‘precharging’ the pixels surrounding the pattern to attract more of the return current [10]. However, such techniques significantly complicate the system design. One issue is that they require a higher duty cycle of the laser illumination during each frame, which carries safety and battery lifetime concerns. Besides, there is not yet a universal algorithm for generating optimal images during the ‘precharging’ phase. Therefore, the optimization described here foregoes such software methods in favor of optimizing only the implant.

Since the stimulation strength and contrast decrease with increasing separation between the electrodes and the target neurons, pillar arrays perform significantly better than the flat ones. Ideally, the pillar height should be customized to match the average thickness of the debris layer in a patient, which can be measured with optical coherence tomography (OCT). However, variation in debris thickness within each retina may result in some differences of the perceptual brightness and contrast over the visual field, as quantified in Tables III and IV.

Pillar electrodes have not been tested in human patients yet, but if they integrate with the retina as well as in recent animal studies [32], then bipolar arrays with 30 µm pixels and pillar electrodes should provide the best average resolution of 42 µm for the whole range of debris variation (*<* 57 µm), and single-pixel resolution with debris *<* 47 µm.

Additionally, as recorded in Tables III and IV, we limited the irradiance to avoid axonal stimulation. However, if clinical testing should show no adverse effects related to direct stimulation of RGCs, the irradiance might be increased up to 3.5 mW*/*mm^2^, with a corresponding improvement in the stimulation strength.

The results presented here assume direct correlation between the electric field and attainable resolution. Primarily, we defined the minimum requirements for resolving an image based on the stimulation thresholds and prosthetic acuity measured in PRIMA patients. However, there might be other factors affecting prosthetic visual perception in patients, such as mental agility and the extent of retinal degeneration. Even though the exact performance may vary from patient to patient, considerations presented in this study still guide towards the electrode geometry that maximizes the stimulus strength, pattern contrast, and selectivity of stimulation.

## V. Conclusion

For subretinal multi-electrode arrays, not only the amount of injected current but also the geometry of active and return electrodes plays a crucial role in the efficacy and selectivity of neural stimulation. Suboptimal designs result in weak stimulation, low contrast, or direct activation of non-target cells. In contrast, this study demonstrates that optimizing the electrodes in subretinal photovoltaic arrays could enhance visual acuity by as much as five times compared to current clinical results. Similar considerations and models can be used for optimizing other electro-neural interfaces. For example, this modeling pipeline allows the assessment of neural stimulation in presence of glial scarring - a common cause of electrode failure. Furthermore, it provides a tool for designing neural prosthetics to be robust within the expected range of uncertainties in tissue thickness and resistivity, which is critical for long-term clinical success.

## Acknowledgment

Nathan Jensen reports employment by Science Corporation. Daniel Palanker reports a consulting relationship with Science Corporation. Daniel Palanker’s patents in the field of retinal prosthesis are owned by Stanford University and licensed to Science Corporation. Other authors declare no competing financial interests that could have appeared to influence the work reported in this paper.

