## Supplemental Information for "Maximizing the fidelity of a photovoltaic subretinal prosthesis for human patients"

### S.I. Optimization Procedures

#### A. Bipolar Electrode Optimization

The following optimization procedure was performed on implants with pixel widths of 20, 30, 40, 50, and 75 µm. All metrics were calculated using a Landolt C projected at 30 Hz, with 9.8 ms pulses at an irradiance of $3\,\mathrm{mW/mm^2}$. The resulting electric potential in the retina is averaged during the illumination pulse. First, the optimal shunt value was found using the method described in the main text (section III-A) of the paper. Next, the pillar height of the active electrodes was swept from 1 to 35 µm. As the electrode height increased, the stimulation strength, contrast, and neural selectivity improved unimodally until the pillar height matched the modelled debris thickness of 35 µm. Therefore, a 35 µm pillar height was selected. Finally, the radius of the active electrode and the width of the return electrode were varied. We used a uniform grid search based on the percentage of the pixel area covered by the two electrodes. The samples were spaced evenly from 5 to 40%, with an increment of 5%, and each possible combination of the active and return electrodes was simulated within that range.

Once the electrode geometry was selected, COMSOL LiveLink for MATLAB was used to generate the electrodes and calculate their elementary electric fields. The results were passed to RPSim, which calculates the electric field in the retina. Further processing is performed to extract the pattern contrast, and the drop in electric potential along bipolar and ganglion cells. Finally, the objective functions were evaluated using a Pareto frontier to select the device with the best performance. This pipeline is fully automated, and, as such, could also be applied to closed-loop optimization methods using gradient descent or Gaussian processes. The optimization loop is depicted in Fig. S1.

Results for 40 µm pixels are shown in Fig. S2: (a) and (b) depict the maximum and average stimulation strength across bipolar cells in a pattern of Landolt C with 48 µm gap. The device geometries that maximize each metric are different, but there are devices that provide good performance in both. Fig. S2(c) shows the maximum drop in electric potential along an RGC axon. The neural selectivity ratio is shown in (d), based on the maximum potential drop across the BCs and RGCs ($\Delta V_{BC}/\Delta V_{RGC}$). Contrast, shown in (e), always remains above the best-case PRIMA performance and, as such, is never a limiting factor for this electrode geometry. The Pareto frontier is shown (f) for the variables of stimulation strength and the neural selectivity ratio.

To select a Pareto optimal point, we have chosen a device with the highest neural selectivity that did not significantly compromise the bipolar cell stimulation strength, which is necessary to ensure good performance. It would be possible to use a more objective metric, such as providing the highest drop in electric potential across the bipolar cells, with an electric field scaled such that the voltage drop across the ganglion cell axons does not exceed their stimulation threshold of 20 mV. Such a device tends to have larger return and smaller active electrodes, resulting in a very strong electric field near the electrode and minimal crosstalk between pixels. However, with thicker debris where the smallest Landolt C resolvable by a certain pixel width cannot reach the stimulation threshold for bipolar cells, the crosstalk would be actually advantageous - allowing larger patterns to add up and exceed the stimulation threshold. As such, for patients with thick debris, thinner return and larger active electrodes provide a smaller resolvable Landolt C than the alternative. With that in mind, we selected the Pareto optimal devices listed in the text.

#### B. Monopolar Electrode Optimization

In a similar manner to adjusting the active and return electrode areas for the bipolar devices, the ratio of the capacitance of the return electrode and total capacitance of the active electrodes can be adjusted for monopolar implants. This can be accomplished using several different methods: 1) Reduction in the width of the global return electrode will increase its access resistance, which results in a larger negative potential at the retinal ganglion cell axons. 2) Use of a material with lower capacitance per unit area than SIROF, such as platinum or even titanium will keep the access resistance the same while lowering the ratio of the return to active electrode capacitance. 3) Moving the return electrode to the back side of the device allows for a wider variety of combinations of electrode widths and specific capacitance per unit area, and also prevents the

Submitted for review. Studies were supported by the National Institutes of Health (R01-EY-035227 and P30-EY-026877) and the Department of Defense (W81XWH-22-1-0933).

1. Department of Electrical Engineering, Stanford University, Stanford, CA 94305 USA
2. Department of Ophthalmology, Stanford University
3. Hansen Experimental Physics Laboratory, Stanford University
4. School of Life Sciences, EPFL, Lausanne, Switzerland

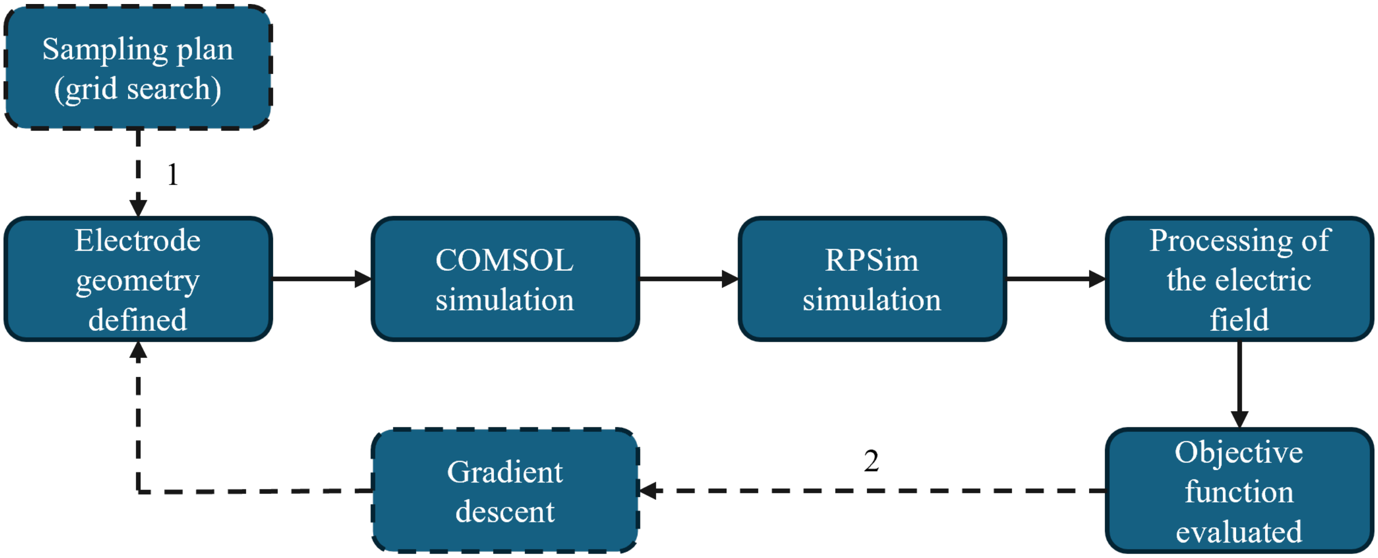


Fig. S1: Flowchart showing the electrode optimization procedure. Dashed paths indicate how this platform can be used in both 1) open loop optimization and 2) closed loop optimization.

negative potential from reaching the RGC axons. Results of such optimization are presented in the main text.

Since contrast is the major limiting factor for monopolar implants, the smallest return/active ratio provides the best performance. In this approach, the current returns through the active electrodes and their shunt resistors in dark pixels rather than the return electrode. However, this method limits the total amount of current that can be injected into the retina. For a uniform beam, this translates to a maximum fraction of the device area that can be illuminated at once (image fill factor), while maintaining the required current injection. For patterns with a small fill factor, i.e. illuminating only a few pixels, the performance will be unaffected, as shown in the top row of Fig. S3 for a single Landolt C. However, full field illumination results in a quick drop of current due to the small return electrode capacitance, as shown in the bottom row of Fig. S3. In this example, the patterns were projected with the same illumination described for the Landolt C above. The capacitance of the large return electrode in this example is 5 times that of all the active electrodes combined, and the small return is 0.01 of the active electrodes' capacitance.

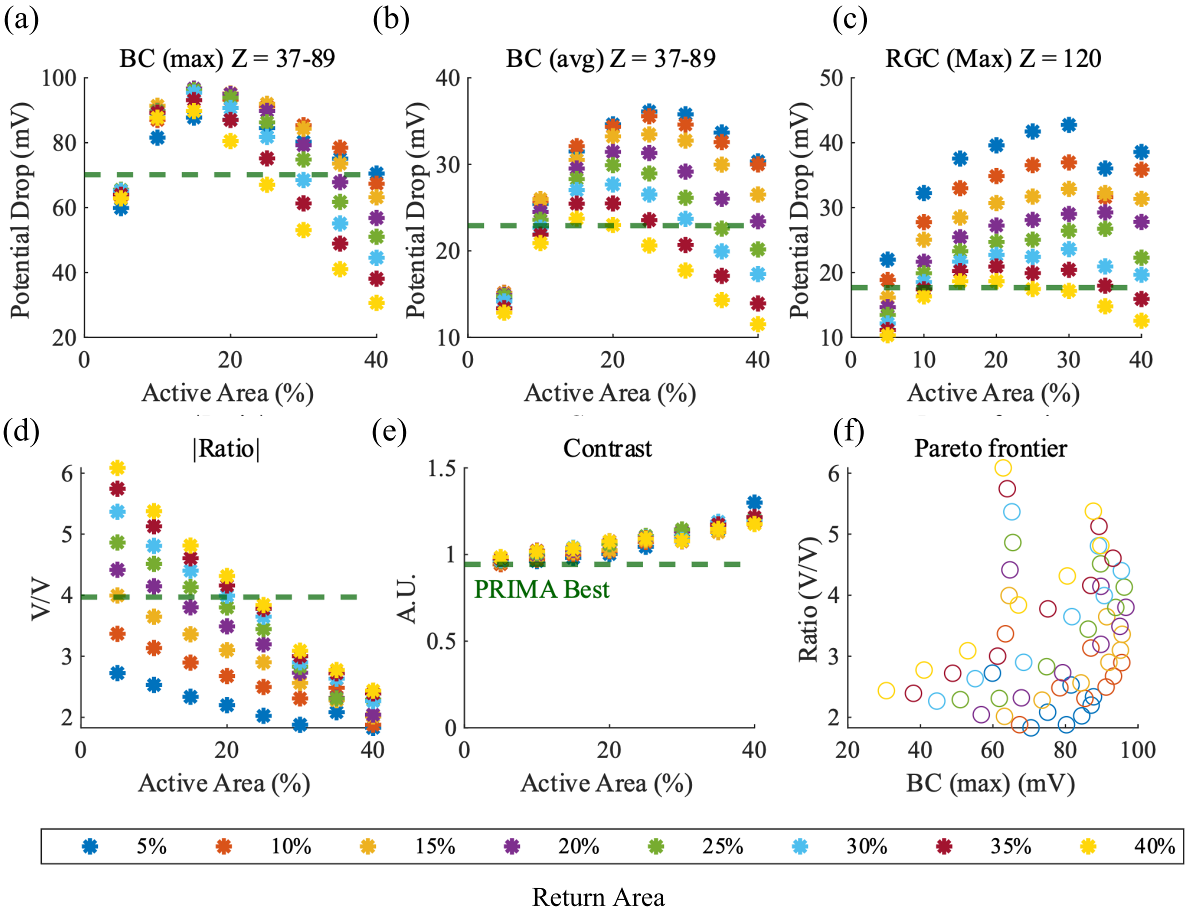


Fig. S2: The performance of all sampled devices with a 40 µm electrode pitch and 35um tall pillars evaluated using a Landolt C with a 48 µm gap width. Horizontal dashed lines indicate the best-case performance of the PRIMA device for a Landolt C of a 120 µm gap width. The horizontal axis depicts the percentage of the device area allocated to the active electrode, and the ratio of the return electrode area to the pixel area is shown using different colors, labelled in the legend. (a) The maximum drop of electric potential across the bipolar cells. (b) The drop of electric potential across the bipolar cells, averaged over the illumination pattern (c) The maximum electric potential drop along the retinal ganglion cell axons (d) The neural selectivity ratio, defined as the ratio of the maximum stimulation strengths of bipolar and ganglion cells (e) The contrast in electrical stimulation between the Landolt C and its gap (f) The pareto plot showing the stimulation strength of bipolar cells and the neural selectivity ratios for every sampled device with 40 µm pixels. Each point corresponds to a different combination of the active and return electrode sizes.

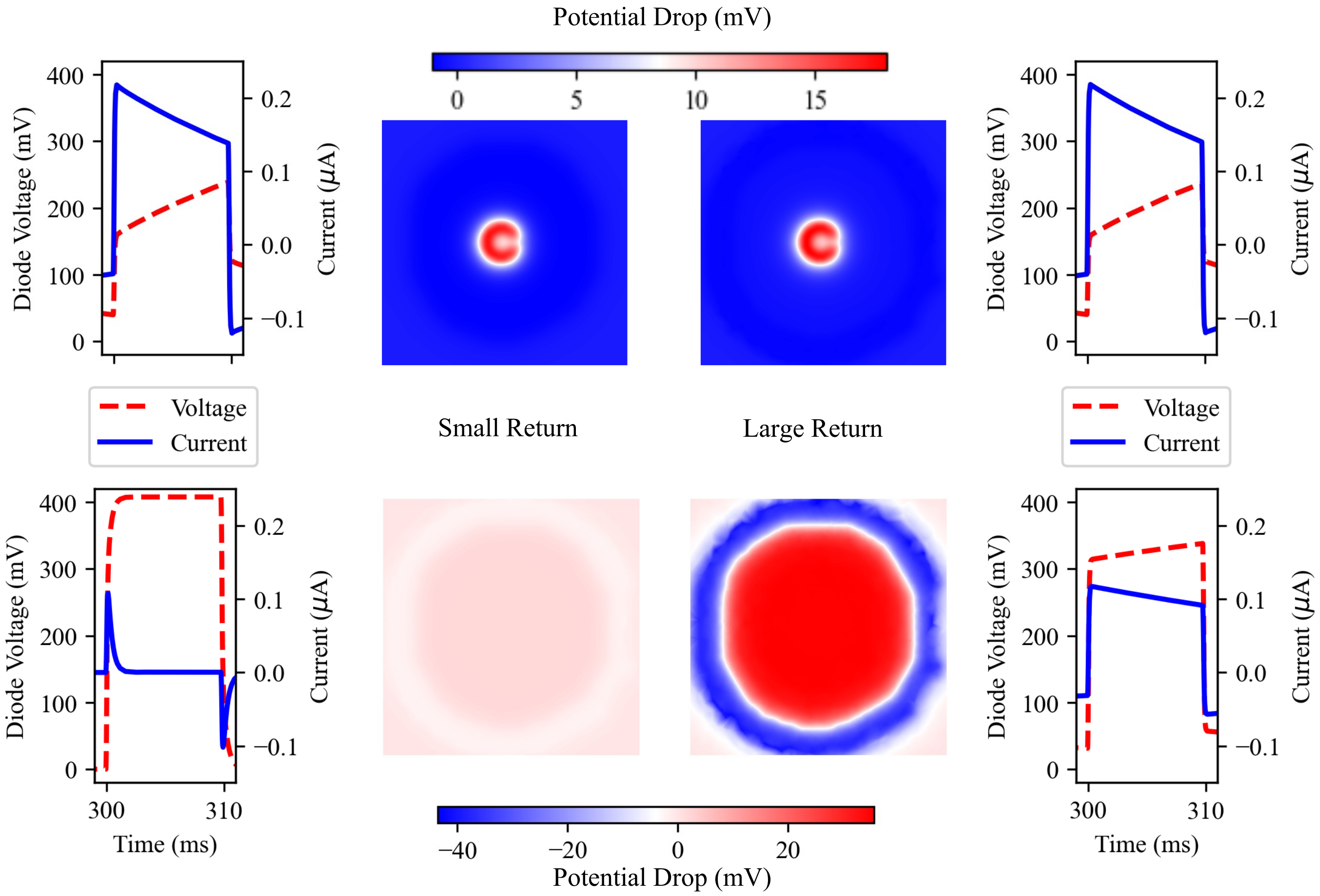


Fig. S3: Drop in electric potential across bipolar cells with a Landolt C of 48 μm gap (top) and a full field illumination (bottom), shown for devices with a small return capacitance (left) and a large one (right). Next to each pattern we show the corresponding current injected into the neural tissue (blue, solid) as well as the voltage on the diodes (red, dash) during the pulse.
